# Point mutations in IMPDH2 which cause early-onset neurodevelopmental disorders disrupt enzyme regulation and filament structure

**DOI:** 10.1101/2023.03.15.532669

**Authors:** Audrey G O’Neill, Anika L Burrell, Michael Zech, Orly Elpeleg, Tamar Harel, Simon Edvardson, Hagar Mor Shaked, Alyssa L Rippert, Tomoki Nomakuchi, Kosuke Izumi, Justin M Kollman

## Abstract

Inosine 5’ monophosphate dehydrogenase (IMPDH) is a critical regulatory enzyme in purine nucleotide biosynthesis that is inhibited by the downstream product GTP. Multiple point mutations in the human isoform IMPDH2 have recently been associated with dystonia and other neurodevelopmental disorders, but the effect of the mutations on enzyme function has not been described. Here, we report identification of two additional affected individuals with missense variants in *IMPDH2* and show that all of the disease-associated mutations disrupt GTP regulation. Cryo-EM structures of one IMPDH2 mutant suggest this regulatory defect arises from a shift in the conformational equilibrium toward a more active state. This structural and functional analysis provides insight into IMPDH2-associated disease mechanisms that point to potential therapeutic approaches and raises new questions about fundamental aspects of IMPDH regulation.

**Significance Statement:** Point mutations in the human enzyme IMPDH2, a critical regulator of nucleotide biosynthesis, are linked to neurodevelopmental disorders, such as dystonia. Here, we report two additional IMPDH2 point mutants associated with similar disorders. We investigate the effects of each mutation on IMPDH2 structure and function *in vitro* and find that all mutations are gain of function, preventing allosteric regulation of IMPDH2 activity. We report high resolution structures of one variant and present a structure-based hypothesis for its dysregulation. This work provides a biochemical basis for understanding diseases caused by *IMPDH2* mutation and lays a foundation for future therapeutic development.

## Introduction

Dystonia is a neurological movement disorder associated with many different genetic variants. Recent studies have identified mutations in IMPDH2, a key regulatory enzyme in purine nucleotide biosynthesis, associated with dystonia and other neurodevelopmental disorders. Purine nucleotides are essential components of cells where they serve as signaling molecules, energy sources, and precursors of RNA and DNA. Purine salvage pathways function alongside *de novo* synthesis pathways to maintain purine pools during steady state, but *de novo* biosynthesis is upregulated during proliferation to meet increased purine demand (1, 2). IMPDH catalyzes the rate-limiting step of *de novo* guanine nucleotide biosynthesis - the conversion of inosine 5’- monophosphate (IMP) to xanthosine 5’-monophosphate (XMP) (3, 4). Because IMP is also a precursor in the *de novo* synthesis of adenine nucleotides, IMPDH sits at a key metabolic branchpoint that controls flux between adenine and guanine nucleotide production (5–7).

To control this branch point, IMPDH is tightly regulated at multiple levels including through allosteric regulation by purine nucleotide binding and reversible assembly into filaments (8–14) (Fig. 1). The IMPDH protomer consists of a catalytic domain and a regulatory Bateman domain, connected by a flexible hinge. IMPDH constitutively assembles tetramers through catalytic domain interactions, and tetramers reversibly dimerize into octamers through interactions of the Bateman domains in response to binding of ATP or GTP in three allosteric sites. Binding of ATP in sites 1 and 2 promotes Bateman domain interactions in an extended, active conformation. Binding of GTP in site 2 and site 3, which is only formed in the compressed conformation, stabilizes a compressed, inactive conformation. Thus, GTP acts as an allosteric inhibitor by controlling the transition from extended to compressed conformations (9).

**Fig. 1.**
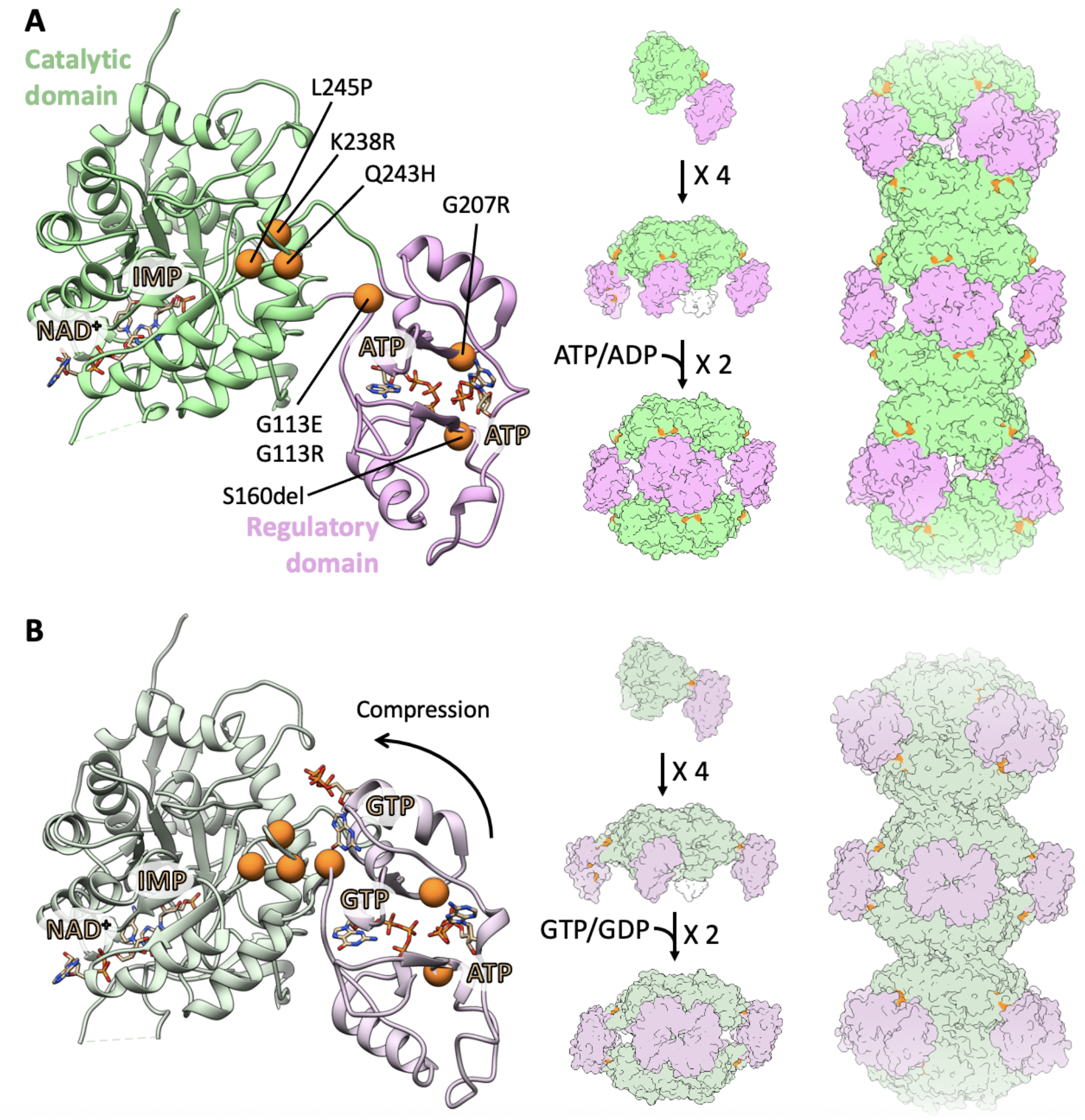
Locations of mutations in IMPDH2. Locations of all seven mutations (orange) are mapped to the IMPDH2 monomer in the extended, active state (A; PDB: 6U8N) and in the compressed, inhibited state (B; PDB: 6U9O). IMPDH2 monomers assemble into tetramers, octamers, and helical filamentous polymers with D4 symmetry.

IMPDH2 is further regulated by assembly into filaments of stacked octamers, which reduces affinity for GTP by disfavoring the compressed conformation (12, 13). Filament formation is induced in cells under conditions with increased demand for guanine nucleotides, such as in response to IMPDH inhibition and starvation (15–22). In the presence of ATP, IMPDH filaments are conformationally heterogeneous, sampling a range of states between symmetrically extended and asymmetrically compressed. In the presence of GTP, the filaments more uniformly adopt a compressed conformation.

In humans, there are two IMPDH isozymes with 84% sequence identity (23). IMPDH1 is constitutively expressed in most cells as a housekeeping gene, while expression levels of IMPDH2 are higher in developing tissues (24, 25). IMPDH2 expression is also selectively enhanced in cancerous cells such as human brain tumors, sarcoma cells, and leukemic cells (24, 26–28). Both isoforms assemble filaments, but for IMPDH2 only, incorporation into filaments reduces sensitivity of the enzyme to GTP inhibition by preventing complete compression of the octamer (13, 14). Assembly of enzymes into filaments is a commonly observed mechanism that cells leverage to regulate metabolic processes (29–31).

Five mutations in *IMPDH2* were recently identified in patients with early-onset neurodevelopmental diseases, including dystonia (32). All of the mutations are located in the regulatory domain or near the hinge that connects the regulatory and catalytic domains (Fig. 1). A series of similar mutations in *IMPDH1* have been associated with retinal degeneration (33– 37), and a subset of these were recently shown to disrupt GTP feedback inhibition (14, 38). The role of IMPDH2 is believed to be essential for cell proliferation and growth, as knockout of IMPDH2 in mice is embryonic lethal (39). Mice deficient in IMPDH1 and heterozygous for IMPDH2 had reduced IMPDH activity, but no developmental defects, supporting the hypothesis that IMPDH dysregulation, rather than a net decrease in its activity, is causative of disease (39, 40). However, more recently, a heterozygous early termination in exon 1 of *IMPDH2* was identified in a patient with dystonia (41), suggesting a decrease in IMPDH2 activity could also be causative of disease.

Here, we report two additional variants, L245P and K238R, of human *IMPDH2* associated with neurodevelopmental disease and show that neurodevelopmental disease-associated point mutations disrupt GTP feedback inhibition. Cryo-EM structures of the L245P mutant show that it can access both extended active and compressed inhibited conformations, but with the equilibrium between these conformational states disrupted relative to wildtype (WT). We propose a mechanism of dysregulation of the L245P variant in which the transition to the inhibited compressed state is disfavored. Other mutations result in a variety of structural phenotypes that suggest different mutations may have different molecular mechanisms of dysregulation.

## Results

### Identification of the L245P variant

The proband (Fig. 2), a 3-year-old female, presented with global developmental delay, congenital anomalies (pulmonic stenosis, hip dysplasia), hypotonia, and dysmorphic features. The torticollis and abnormal posturing may be a form of dystonia in this patient. The full clinical description of the patient is reported in Supplemental Notes.

**Fig. 2.**
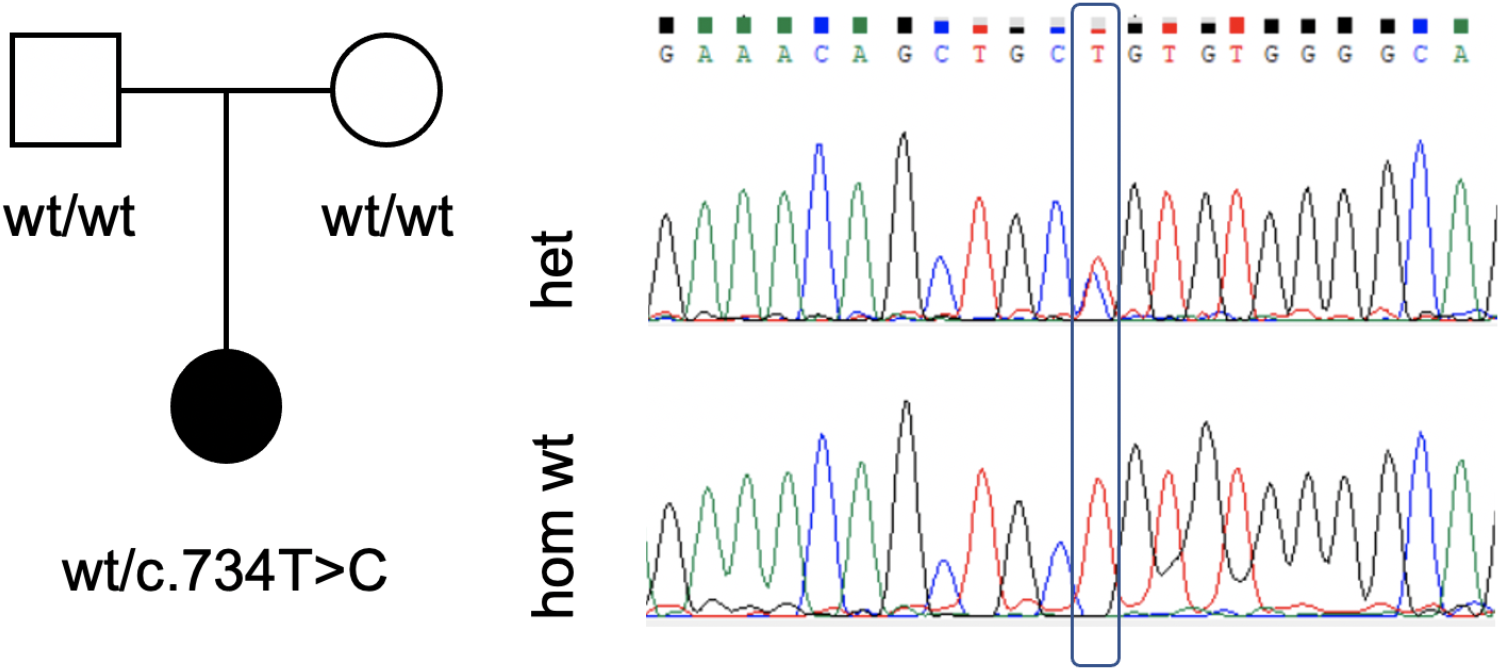
Genetic evaluation of the L245P variant. Pedigree of the proband with the *de novo* L245P variant of *IMPDH2* (left). Male individuals are represented by squares, and female individuals are represented by circles. Sanger sequencing confirmed the heterozygous c.734T>C mutation (right).

Genetic evaluation in the proband included a chromosomal microarray (CMA) which did not identify any pathogenic or likely pathogenic genomic deletions or duplications, followed by trio exome sequencing. A de novo heterozygous variant was identified in IMPDH2: chr3:49064205[hg19]; NM_000884.3; c.734T>C; p.(Leu245Pro). This was confirmed by Sanger sequencing (Fig. 2). The variant is rare (not found in gnomAD or in the local database of ∼15,000 exomes), alters a conserved amino acid (GERP 5.97), has a high CADD score of 29.7, and is predicted to be damaging by multiple bioinformatic algorithms (MutationTaster, SIFT, Revel, and others). In addition, a de novo heterozygous variant was identified in LRP1: chr12:57589465[hg19]; c.8463dup; p.(Glu2822Ter). Pathogenic variants in LRP1 have recently been associated with developmental dysplasia of the hip (42) and this variant therefore may have contributed to this phenotype in the proband.

### Identification of the K238R variant

The proband is a now 3-year-old male who presented to genetics evaluation at 9 months of age due to hypotonia and global developmental delays, including gross motor and speech delay. The full clinical description of the patient is reported in the Supplemental Notes. Photos of the proband at different ages are shown in Supplemental Fig. 1.

Chromosomal SNP microarray, Prader-Willi/Angelman syndrome methylation analysis, and metabolic screening labs were all performed and non-diagnostic. Exome sequencing was then recommended at 16 months of age and identified a de novo, likely pathogenic variant in IMPDH2 (c.713A>G, p.Lys238Arg).

### IMPDH2 variants are defective in GTP regulation

To determine whether disease associated *IMPDH2* mutations have a direct effect on enzyme activity or regulation, we assayed purified recombinant enzymes *in vitro*. We observed modest variation in the apparent V_max_ of the enzymes and K_0.5_ values for IMP and NAD+ among the IMPDH2 variants relative to WT enzyme, suggesting that basal activity is not severely affected by the mutations (Table 1). We next tested whether the disease mutations affect GTP inhibition (Fig. 3). WT IMPDH2 filaments are inhibited by GTP with an IC50 of 577 μM under our assay conditions (Fig. 3); as previously reported, we find that IMPDH2 retains a basal level of activity even in the presence of saturating GTP (12, 13). Each mutant we tested retains significant activity up to 5 mM GTP (Fig. 3). While L245P, one of the mutations reported here, could be inhibited at much higher GTP concentrations, with an estimated IC50 of 7 mM, this value is far above the usual physiological concentration range of GTP (Supplemental Fig. 2). Thus, each neurodevelopmental mutant dramatically compromises feedback inhibition by GTP, suggesting that disease phenotypes may be related to hyperactivity of IMPDH2 under conditions in which the WT enzyme would be inhibited.

**Table 1:** Kinetic parameters of IMPDH2 variants compared to WT IMPDH2.

| Variant | IMP | | NAD<sup>+</sup> | | V<sub>max app</sub><br>($\mu\text{M NADH min}^{-1}$) |
| --- | --- | --- | --- | --- | --- |
| | K<sub>0.5</sub> ($\mu\text{M}$) | Hill | K<sub>0.5</sub> ($\mu\text{M}$) | Hill | |
| WT | 45.6 | 7.4 | 44.9 | 5.3 | 2.2 ± 0.1 |
| G113E | 17.6 | 2.3 | 44.9 | 2.8 | 2.4 ± 0.8 |
| G113R | 20.1 | 2.8 | 31.5 | 3.3 | 2.0 ± 0.5 |
| G207R | 35.5 | 2.8 | 37.1 | 5.8 | 2.1 ± 0.1 |
| S160del | 15.1 | 2.8 | 25.8 | 5.7 | 1.7 ± 0.2 |
| Q243H | 52.1 | 4.9 | 60.2 | 3.0 | 3.6 ± 0.5 |
| L245P | 16.9 | 1.7 | 24.6 | 3.5 | 2.8 ± 0.4 |
| K238R | 17.2 | 2.5 | 33.4 | 9.0 | 1.4 ± 0.1 |

**Fig. 3.**
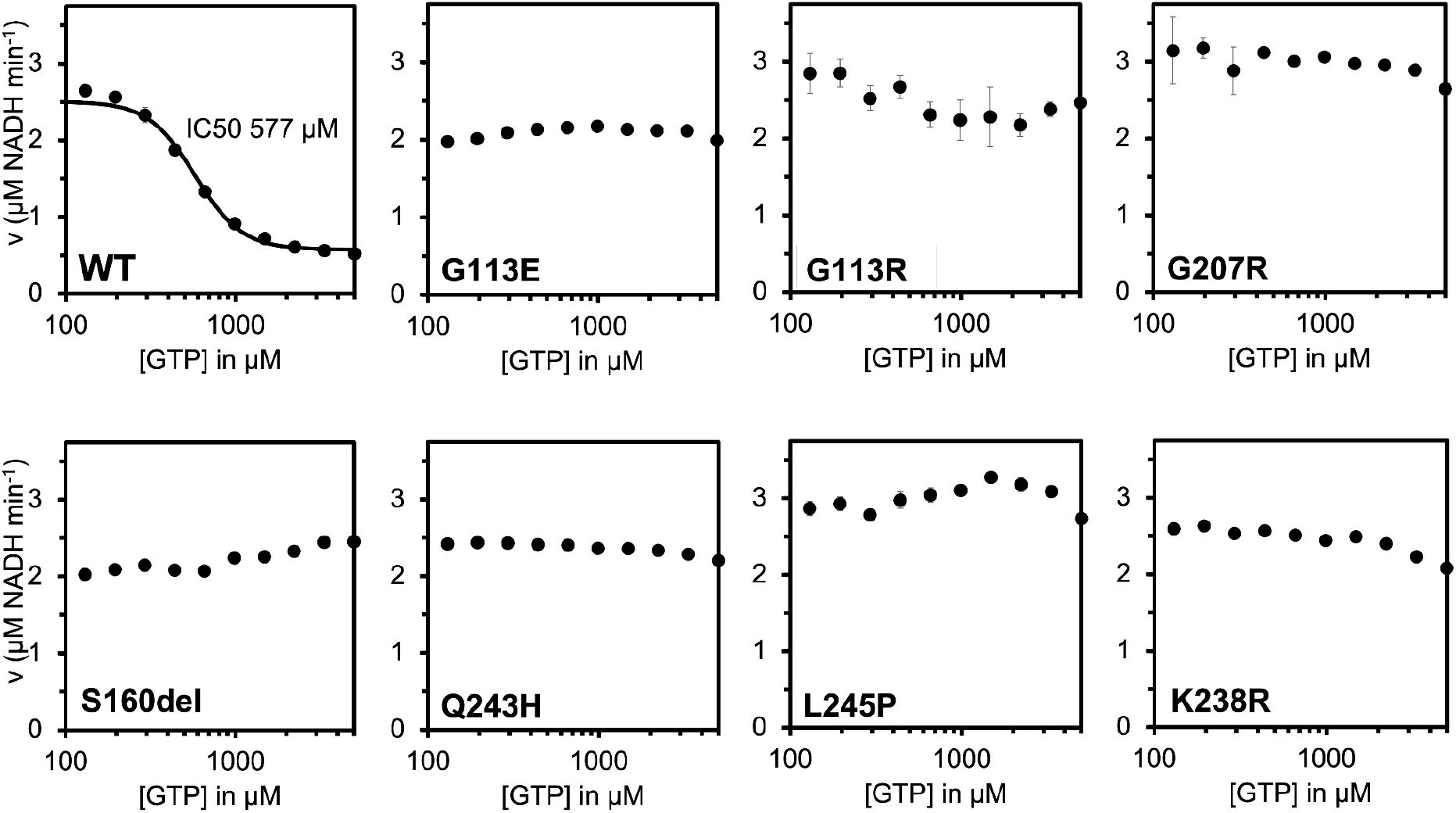
IMPDH2 neurodevelopmental variants disrupt GTP inhibition. GTP inhibition of WT IMPDH2 and IMPDH2 neurodevelopmental variants. Each data point represents the average initial rate of three reactions. Error bars represent standard deviation for n=3 technical replicates. Velocities were calculated from the change in absorbance at 340 nm. Reactions were initiated with 300 μM NAD+ and contained 1 μM enzyme, 1 mM ATP, 1 mM IMP, 1 mM MgCl_2_ and varying concentrations of GTP.

### Structural phenotypes of IMPDH2 mutants

The IMPDH2 neurodevelopmental mutations are in or near the regulatory domain, which controls the extended-compressed structural transition (Fig. 1). This led us to hypothesize that the loss of GTP regulation we observe arises from an inability to transition into the compressed inhibited conformation. To assess changes to the structure and conformation in IMPDH2 mutants, we used negative stain electron microscopy. Under activating and inhibiting conditions for the WT enzyme, it is relatively straightforward to assess whether the enzyme is extended or compressed by directly observing the helical rise of filaments (12–14). Because the mutants have normal basal activity but are not inhibited by GTP, we anticipated that in the absence of GTP, the mutants should resemble the WT in the extended, active conformation. In the presence of GTP, which causes compression of WT filaments, we predicted we would not observe compression. However, negative stain analysis of IMPDH2 revealed surprising large-scale structural differences among the mutants (Fig. 4, Table 2).

**Table 2:** Summary of negative stain results from Figure 4.

| **Variant** | **ATP** | **GTP** |
| --- | --- | --- |
| WT | Extended filaments | Compressed filaments |
| G113E | Compressed filaments | Compressed filaments |
| G113R | Compressed filaments | Compressed filaments |
| G207R | Extended filaments | Compressed filaments |
| S160del | No filaments | Octamers |
| Q243H | Extended filaments | Compressed filaments |
| L245P | Extended filaments | Compressed filaments |
| K238R | Extended filaments | Compressed filaments |

**Fig. 4.**
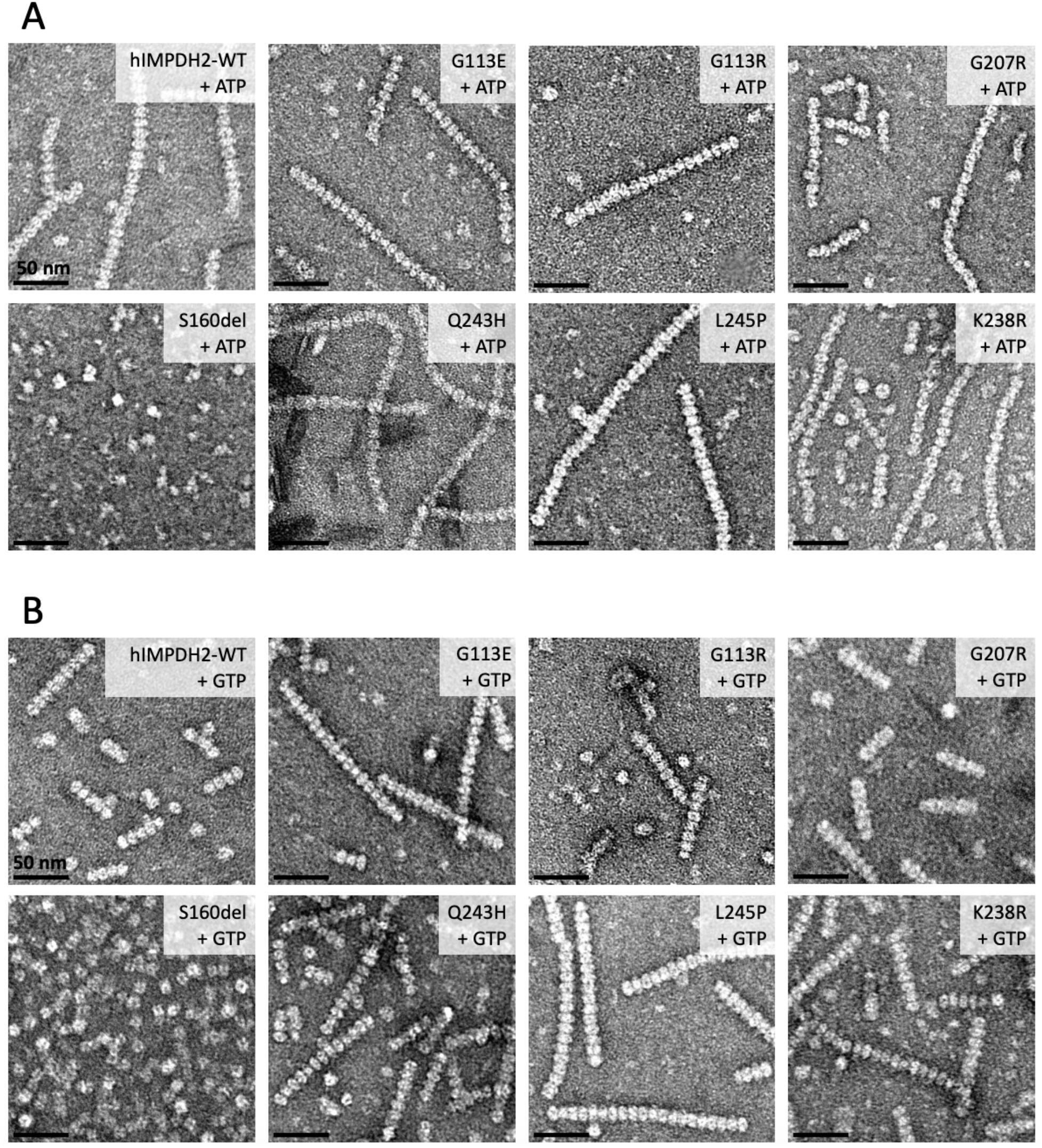
Negative stain EM reveals low-resolution differences between some mutants. Representative negative stain images of 2 μM enzyme with either 1 mM ATP and 1 mM MgCl_2_ (A) or 5 mM GTP (B). The oligomeric and conformational states of some mutants vary from WT. S160del does not form filaments in either condition. G113E and G113R form compressed filaments in the absence of GTP.

First, we examined IMPDH2 mutant structures in the absence of GTP, where all mutants retain WT activity (Fig. 4A). Under this condition, G207R, Q243H, L245P, and K238R closely resemble WT in the canonical extended and active conformations. However, S160del did not assemble filaments at all, instead forming mostly tetramers in the presence of ATP. Surprisingly, both G113E and G113R are uniformly compressed, which has not previously been observed for IMPDH2 in the absence of GTP (9–13). More surprising still, in the presence of GTP, all of the disease mutants are able to assemble filaments in the compressed conformation, with the exception of S160del which forms only octamers, although it was not possible to directly measure whether GTP octamer are in the extended or compressed conformation (Fig. 4B). In all prior studies, a compressed conformation of IMPDH is associated with inhibition (9, 11–13), but the variants described here retain catalytic activity in the presence of GTP, suggesting that additional factors beyond compression must be required for IMPDH2 inhibition (Table 2).

### Filament assembly reduces sensitivity of variants to GTP

We next investigated the effect of filament assembly on the variants’ sensitivity to GTP. The engineered mutation Y12A at the filament assembly interface of IMPDH2 disrupts polymerization, and increases sensitivity of IMPDH2 to GTP inhibition compared to the WT (12, 13). We introduced the Y12A mutation into each of the IMPDH2 mutants and confirmed with negative stain that the double mutants do not form filaments (Supplemental Fig. 3). Next, we performed GTP inhibition assays on the double mutants. Because the S160del mutant does not form filaments at all, we anticipated that the Y12A mutation would not affect enzyme activity (Fig. 4). The S160del+Y12A mutant displayed no significant decrease in activity in the presence of GTP, as expected (Fig. 5). The other six double mutants display some inhibition at high GTP concentrations, with L245P+Y12A being the most sensitive (IC50 = 1.3 mM), followed by K238R+Y12A (IC50 = 1.9 mM). None of the double mutants were as sensitive to GTP inhibition as the non-assembly mutant alone (IC50 = 200 μM). This suggests that the mutations affect the octameric form of the enzyme as well, and assembly of the variants into filaments would further exacerbate downstream regulatory defects in the cell.

**Fig. 5.**
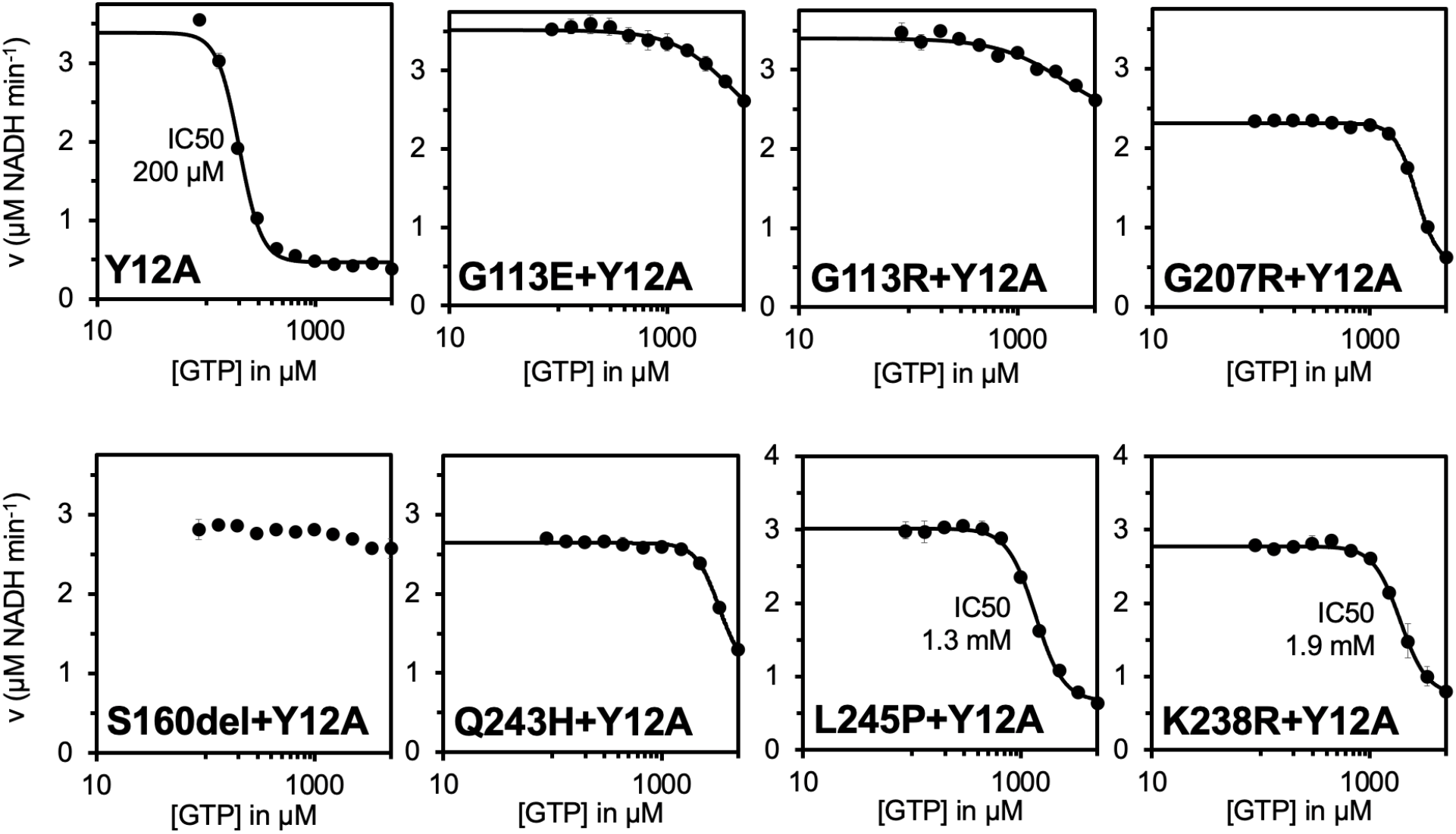
Mutations also disrupt GTP inhibition of free octamers. GTP inhibition curves of Y12A non-assembly mutant and Y12A/disease mutant double mutants of IMPDH2. Each data point represents the average initial rate of three reactions. Error bars represent standard deviation for n=3 technical replicates. Velocities were calculated from the change in absorbance at 340 nm. Reactions were initiated with 300 μM NAD+ and contained 1 μM enzyme, 1 mM ATP, 1 mM IMP, 1 mM MgCl_2_ and varying concentrations of GTP.

### L245P filament structures reveal differences in flexibility

To investigate the structural basis for IMPDH2 dysregulation by neurodevelopmental mutations, we determined structures of L245P in the presence of ATP and GTP. We chose L245P as representative of the most common structural phenotype we observed, with enzymes that undergo the extended to compressed transition in the presence of GTP despite retaining full activity (G207R, Q243H, L245P, and K238R).

First, we determined the structure of L245P in the catalytically active extended conformation, in the absence of GTP. Flexibility in WT IMPDH2 filaments under these conditions arises from heterogeneity in the extended and compressed conformations of protomers within each octamer, which can limit the resolution of cryo-EM structures. We previously developed an image processing strategy for the very flexible WT IMPDH2 filaments, which allows us to computationally separate uniformly extended from partially compressed (bent) filament segments (13). The approach also allows us to generate focused reconstructions of the repeating octameric subunit and the filament assembly interface. Following symmetry expansion and focused classification, we found that the majority of L245P segments were sorted into fully extended classes (73%), compared to only 17% fully extended WT IMPDH2 segments under these conditions (Supplemental Fig. 4) (13). Thus, it appears that L245P reduces octamer flexibility in the active state, leading to a more uniformly extended structure than the very heterogeneous WT filaments.

The extended L245P structures refined to global resolutions that were significantly higher than our earlier WT structures: 2.0 Å for the filament assembly interface and the catalytic core, and 2.6 Å resolution for the octamer-centered reconstruction (Fig. 6C, 6B). The mutation of L245 to proline is clear in the cryo-EM map and does not appear to perturb the conformation of the backbone around the mutation site relative to WT (Fig. 6E). IMP and NAD+ are clearly resolved in the active site, and ATP is well resolved in sites 1 and 2 in the regulatory domain (Supplemental Fig. 5). Overall, the structures are nearly identical to the WT structure under this condition at the level of individual protomers, octamers, and the filament assembly interface (Fig. 6D). Some minor differences in loops of the regulatory domain likely reflect improved accuracy in model building at the improved resolution of the current reconstructions.

**Fig. 6.**
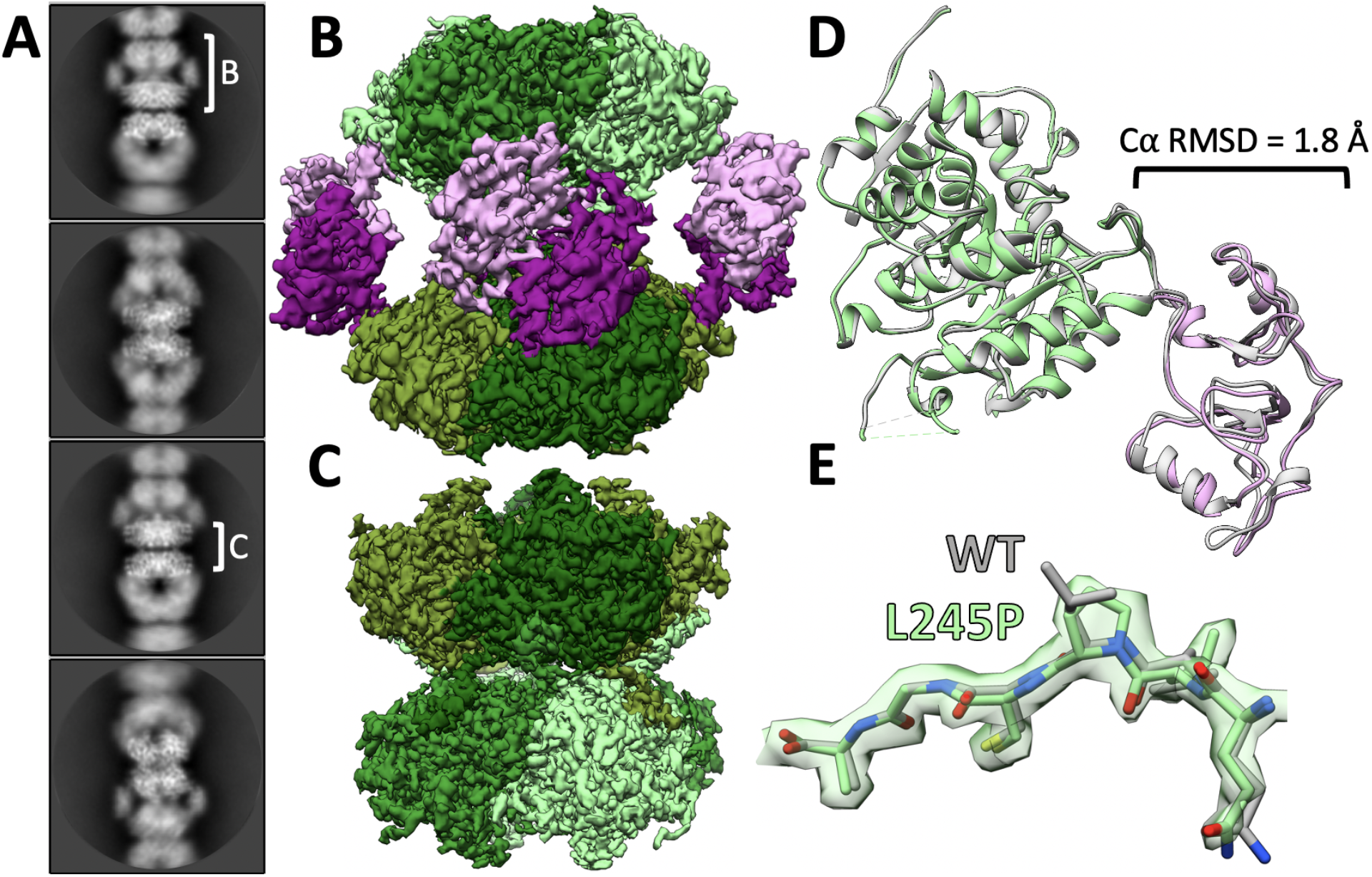
L245P mutant adopts similar extended conformation as the WT. Representative 2D class averages of L245P filaments in the presence of 1 mM ATP, 3 mM IMP, 5 mM NAD+, and 1 mM MgCl_2_ (A). White brackets illustrate the regions on the filament where refinement was focused to produce reconstructions shown in panels B and C. Final cryo-EM reconstructions of octamer-centered (B) and interface-centered (C) filament segments. Regulatory domains are colored in shades of pink, and catalytic domains are colored in shades of green. Alignment of the L245P octamer-centered ribbon model (color; PDB 8G8F) to the WT ribbon model (gray; PDB 6U8N) at the catalytic domain (green). Calculation of C*α* RMSD at the regulatory domain (pink) shows minor differences in the structure of the monomer (D). The density around the mutation site shows that the backbone structure is not affected (E).

Next, we determined the structure of L245P in the presence of 20 mM GTP. At this high GTP concentration, we had previously observed that the WT IMPDH2 forms uniformly compressed filaments, with no significant population of extended or bent octamers (13). However, from initial two-dimensional classification of L245P helical segments, it was clear that the mutant filaments were very heterogeneous (Fig. 7A). The data processing approach described above allowed us to separate out asymmetric bent segments, which closely resemble bent segments of WT IMPDH2 observed only in the absence of GTP (13). The remaining symmetrically compressed filament segments comprise only about 13% of the data set (Supplemental Fig. 6). Compared to the uniformly compressed WT, then, one consequence of the L245P mutation is to increase the heterogeneity of IMPDH2 filaments in the presence of GTP.

**Fig. 7.**
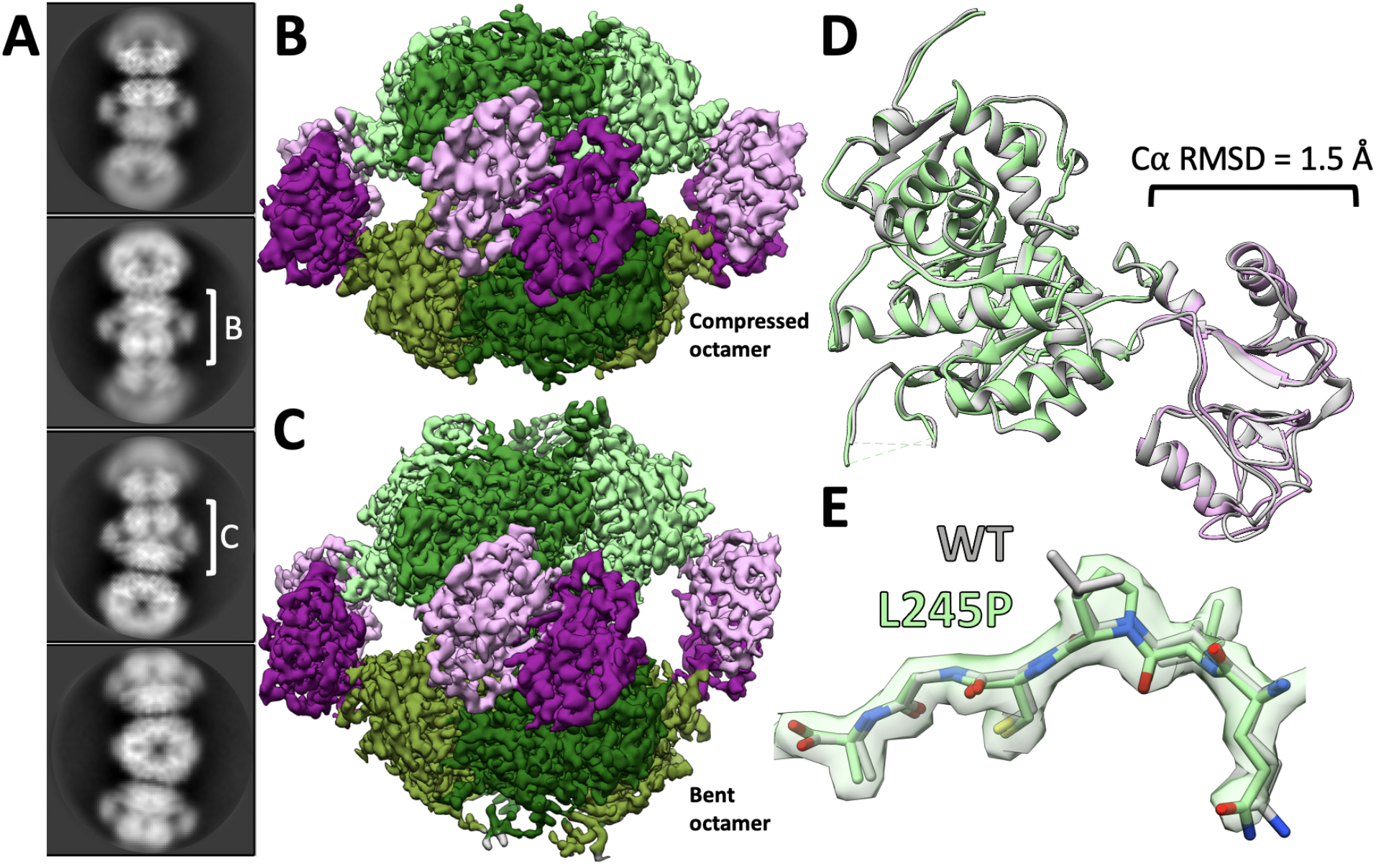
L245P filament is flexible in the presence of GTP. Representative 2D class averages of L245P filaments in the presence of 20 mM GTP, 1 mM ATP, 3 mM IMP, 5 mM NAD+, and 1 mM MgCl_2_ (A). White brackets illustrate the regions on the filament where refinement was focused to produce reconstructions shown in panels B and C. Final cryo-EM reconstructions of straight octamer-centered (B) and bent octamer-centered (C) filament segments. Regulatory domains are colored in shades of pink, and catalytic domains are colored in shades of green. The filament assembly interface reconstruction is shown in Supplemental Figure 7. A model was built into the symmetrically compressed octamer reconstruction. Alignment of the L245P straight octamer-centered ribbon model (color; PDB 8G9B) to the WT ribbon model (gray; PDB 6U9O) at the catalytic domain (green). Calculation of C*α* RMSD at the regulatory domain (pink) again shows minor differences in the structure of the monomer (D). The density around the mutation site shows that the backbone structure is not affected (E).

We used the symmetrically compressed filament segments to determine structures with global resolutions of 2.1 Å for the filament assembly interface and 3.0 Å for the octamer-centered reconstruction (Supplemental Fig. 6). We also determined a structure of the asymmetric, bent filament segment at a global resolution of 2.7 Å (Supplemental Fig. 6). Again, all ligands including GTP were well resolved (Supplemental Fig. 5), and the filament interface was nearly identical to WT. The L245P protomer is also nearly identical to the WT protomer under this ligand condition (Fig. 7D). Thus, IMPDH2-L245P can adopt a canonical compressed structure in the presence of GTP. However, under conditions that support uniform compression and inhibition of the WT enzyme, the mutant remains in an ensemble of partially compressed states.

## Discussion

Maintenance of purine pools is critical in the central nervous system, where purine-based nucleotides and nucleosides have additional functions as second messengers, neurotransmitters, neuromodulators, and trophic agents (43–46). Purine nucleotide and nucleoside pools can be maintained by either salvage pathways or *de novo* pathways, but brain tissue relies primarily on salvage over *de novo* biosynthetic pathways (47). Genetic deficiencies in salvage pathway enzymes result in increased flux through the de novo purine synthesis, leading to the neurodevelopmental disorder Lesch-Nyhan syndrome (48–51). The phenotype of this syndrome, including variably expressed generalized dystonia, motor disability, and cognitive disability, resembles phenotypes in cases of IMPDH2 variants (32). In this study, we show that these IMPDH2 variants all display insensitivity to allosteric inhibition by GTP, supporting the hypothesis that increased activity in the *de novo* purine biosynthetic pathway and subsequent perturbation of purine pools may lead to neurodevelopmental phenotypes.

Like IMPDH2, mutations in the gene for the other human isozyme, *IMPDH1*, also cause disease, in this case the retinal diseases Leber congenital amaurosis and retinitis pigmentosa (33–37). Like the IMPDH2 mutations characterized here, a subset of IMPDH1 mutations cluster near the allosteric domain or inter-domain hinge, and disrupt GTP inhibition (9, 14). Importantly, four of the five IMPDH1 mutations that disrupt GTP regulation prevent IMPDH1 from adopting the compressed conformation, which we proposed as the mechanism of GTP dysregulation. We anticipated that the IMPDH2 mutations studied here would have the same effect but were surprised to see that in each case we could measure, GTP appeared to cause compression of IMPDH2 in low-resolution negative stain micrographs (Fig. 4). Because these enzymes all retain high activity levels in the presence of GTP, this result suggests that contrary to prior models, compression of IMPDH2 alone is not sufficient for allosteric inhibition.

IMPDH2-L245P structures provide insight into the seeming contradiction of the enzyme retaining WT activity in a compressed conformation. We found that the canonical extended and compressed states are virtually identical to the WT structures under the same conditions, including the binding of GTP to inhibitory allosteric sites. However, the conformational variability of filaments in different ligand states varies significantly. Active WT IMPDH2 is in an ensemble of structural states with different numbers of compressed and extended protomers within octamers giving rise to multiple bent conformations, and only a minority (17%) of octameric filament segments were classified as symmetrically extended (13). By comparison, the vast majority (73%) of L245P octamers are symmetrically extended, suggesting that one effect of the mutation is to rigidify the enzyme in the extended state. Conversely, under high concentrations of GTP, WT IMPDH2 octamers are uniformly compressed, while only a minority of L245P octamers (13%) are fully compressed, the majority being in an ensemble of bent conformational states that resemble the active WT ensemble. That is, what appeared in low resolution negative stain micrographs to be the compressed conformation was revealed to be an ensemble of partially compressed conformations using higher resolution cryo-EM approaches. Thus, L245P appears to disrupt GTP regulation by shifting the conformational equilibrium away from the inhibited, compressed conformation (Fig. 8).

**Fig. 8.**
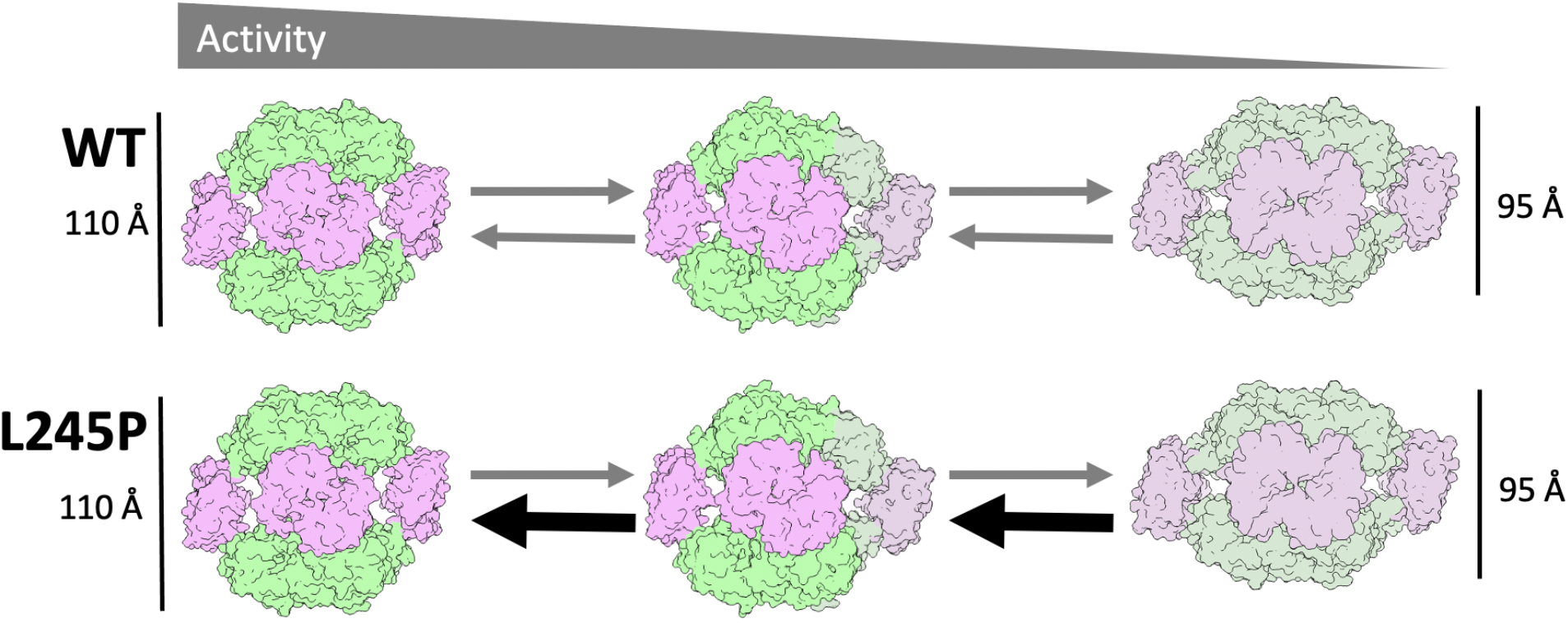
Model for the effect of L245P on IMPDH2 conformational equilibrium. WT IMPDH2 samples conformational space between fully extended and fully compressed octamers. L245P shifts that equilibrium to favor extended and bent conformations, resisting complete compression and maintaining activity.

Whether a disrupted conformational equilibrium underlies the GTP regulation defect for other IMPDH2 disease mutants remains to be determined, although our results suggest that some of the point mutations may work through different mechanisms. For example, S160del was the only mutation that disrupts filament assembly, which may play a role in its dysregulation. Because the seven mutations do not all lead to the same structural phenotypes at the level of polymerization or conformational state (Fig. 4), it will be important to characterize each one to elucidate mechanisms of disease for each. Understanding individual mechanisms of dysregulation may also provide insight into variation in the onset and severity of disease phenotypes.

Additionally, two mutations at residue 113 appear to cause IMPDH2 to be constitutively in a compressed conformation in the absence of GTP. These mutations raise the question of how compressed IMPDH2 retains full WT levels of activity and suggests that compression alone is not sufficient to inactivate the enzyme. The G113E and G113R variants, therefore, may serve as a useful tool for dissecting previously unappreciated mechanisms of regulation of IMPDH2.

Assembly of IMPDH2 into filaments exacerbates the reduction in GTP sensitivity of the disease mutations. We have previously established the role for polymerization in reducing sensitivity to GTP inhibition, which increases the working range of the enzyme under conditions of elevated purine nucleotide demand (13, 14). When filament assembly was blocked, all of the point mutants partially restore sensitivity to GTP inhibition, although not to WT levels (Fig. 5). The sensitivity to GTP inhibition of S160del, which itself prevents filament assembly, was not restored by introduction of our engineered non-assembly mutation Y12A.

Our results suggest potential therapeutic approaches that might be considered for treating patients with IMPDH2-associated developmental defects. The mutations are gain of function, allowing high levels of IMPDH activity at otherwise inhibitory concentrations of GTP. One approach might be the use of known IMPDH inhibitors, to reduce the activity of the enzyme (52–54). Alternative approaches that target IMPDH2 assembly into filaments might also prove effective in some cases; for example disrupting IMPDH2-L245P filament assembly reduces the IC50 for GTP into a more physiologically relevant range and might reduce activity sufficiently to be therapeutically useful. Future work investigating the treatment of IMPDH2 associated defects using inhibitors in cell-based systems and model organisms may prove fruitful.

In both IMPDH1 and IMPDH2, disease-linked mutations in and around the Bateman domain result in the same type of biochemical defect affecting the nervous system. In the case of IMPDH1, this defect results in the degeneration of photoreceptors, possibly due to their unique dependence on IMPDH1 for ATP and cGMP (55–59). Imbalance of nucleotide pools in photoreceptors leads to photoreceptor death (60–62). In the case of IMPDH2, the defect results in neurodevelopmental disorders. In both cases, the dysregulation of IMPDH disrupts the delicate balance of purine pools in the nervous system.

## Materials & Methods

### Exome sequencing and Sanger validation of the IMPDH2 variants

After informed consent, genomic DNA was extracted from peripheral blood samples of the proband with the L245P variant and both parents. Exonic sequences were enriched in the DNA sample using the IDT xGen Exome Research Panel V2.0 capture combined with xGen Human mtDNA Research Panel v1.0 (Integrated DNA Technologies, Iowa, United States), and sequenced on a NovaSeq 6000 sequencing system (Illumina, San Diego, CA) as 100-bp paired-end runs. Data analysis including read alignment and variant calling was performed with DNAnexus software (Palo Alto, CA) using default parameters, with the human genome assembly hg19/GRCh37 as reference. Variants were filtered out if they were off-target (intronic variants >8bp from splice junction), synonymous (unless <4bp from the splice site), or had minor allele frequency (MAF) >0.01 in the Genome Aggregation Database (gnomAD) or in our in-house exome database. The IMPDH2 and LRP1 variants were confirmed by Sanger sequencing in the affected individual, and were not found in either parent. This work was done at Hadassah Medical center, Jerusalem, Israel.

Genetic testing of the individual with the K238R variant was done as clinical trio exome sequencing through Invitae. This work was done at the Children’s Hospital of Philadelphia.

### Recombinant IMPDH expression and purification

Purified IMPDH protein was prepared as described previously (12–14). Briefly, IMPDH2 variants were cloned into a pSMT3-Kan vector with an N-terminal 6xHis-SMT3/SUMO tag. Constructs were transformed into BL21 (DE3) *E. coli* and cultured in LB at 37°C to an OD600 of 0.9. Overexpression was induced with 1mM IPTG for 4 hr at 30°C. Cells were collected by centrifugation. Cell pellets were resuspended at 4°C in lysis buffer (50 mM KPO_4_, 300 mM KCl, 10 mM imidazole, 800 mM urea, pH 8) with a dounce homogenizer, and the cells were lysed with an Emulsiflex-05 homogenizer. Lysate was cleared by centrifugation, and 6xHis-SMT3/SUMO tagged IMPDH2 was initially purified by Ni affinity chromatography using either a HisTrap FF column (GE Healthcare Life Sciences) on an Äkta Start chromatography system or a handpacked HisPur™ Ni-NTA resin (Thermo Scientific), eluting with 50 mM KPO_4_, 300 mM KCl, 500 mM imidazole, pH 8. Fractions containing IMPDH2 were treated with 1 mg ULP1 protease (63) per 100 mg IMPDH for 1 hour at 4°C to cleave the 6xHis-SMT3/SUMO tag. Following cleavage, 1 mM dithiothreitol (DTT) and 800 mM urea were added to inhibit polymerization. Protein was concentrated using a 30,000 MWCO Amicon filter and applied to a Superose 6 column pre-equilibrated in gel filtration buffer (20 mM HEPES, 100 mM KCl, 800 mM urea, 1 mM DTT, pH 8) using an Äkta Pure FPLC system. Peak fractions were concentrated using a 10,000 MWCO Amicon filter, flash-frozen in liquid nitrogen, and stored in single use aliquots at −80°C.

### IMPDH activity assays

Protein aliquots were diluted in assay buffer (20 mM HEPES, 100 mM KCl, 1 mM DTT, pH 7.0) and pre-treated with varying concentrations of ATP, GTP, and IMP for 15 minutes at 25°C in 96 well UV half-area transparent plates (Corning model 3679). Reactions (100 μL total) were initiated by addition of varying concentrations of NAD+. NADH production was measured over time in increments of 1 minute for 15 minutes by absorbance at 340 nm using a Varioskan Lux microplate reader (Thermo Scientific) at 25°C. Absorbance was correlated with NADH concentration using a standard curve. Specific activity was calculated by linear interpretation of the reaction slope for a 4 minute window beginning 1 minute after reaction initiation. All data points reported are an average of 3 measurements from the same protein preparation. Error bars are standard deviation. Fits for activity assays were calculated using the Hill-Langmuir equation *V* = *V*_*max*_ + [*S*]^*n*^/((*K*0.5)^*n*^ + [*S*]^*n*^) and IC_50_ was calculated using a modified Hill equation *V* = *V*_*min*_ + (*V*_*max*_ − *V*_*min*_)/(1 + (*I*/*IC*_50_)^*hill*^(64).

### Negatively stained electron microscopy

Samples were applied to glow-discharged continuous carbon EM grids and negatively stained with 2% uranyl formate. Grids were imaged by transmission electron microscopy using an FEI Morgagni at 100kV acceleration voltage and a Gatan Orius CCD. Micrographs were collected at a nominal 22,000x magnification (pixel size 3.9 Å).

### Electron cryo-microscopy sample preparation and data collection

Samples were applied to glow-discharged C-flat holey carbon EM grids (Protochips), blotted, and plunge-frozen in liquid ethane using a Vitrobot plunging apparatus (FEI) at 4°C, 100% relative humidity. High-throughput data collection was performed using an FEI Titan Krios transmission electron microscope operating at 300 kV (equipped with a Gatan image filter (GIF) and post-GIF Gatan K3 Summit direct electron detector) using the Leginon software package (65).

### Electron cryo-microscopy image processing

Data collection parameters are summarized in Supplemental Table 3. Movies were collected in super-resolution mode using Leginon, then aligned and corrected for beam-induced motion using Motioncor2, with 2x Fourier binning and dose compensation applied during motion correction (65, 66). CTF was estimated using CTFFIND4 (67). Relion 3.1 was used for all subsequent image processing (68, 69). Each dataset was individually processed but using approximately the same previously published pipeline (13, 14).

**Table 3:**
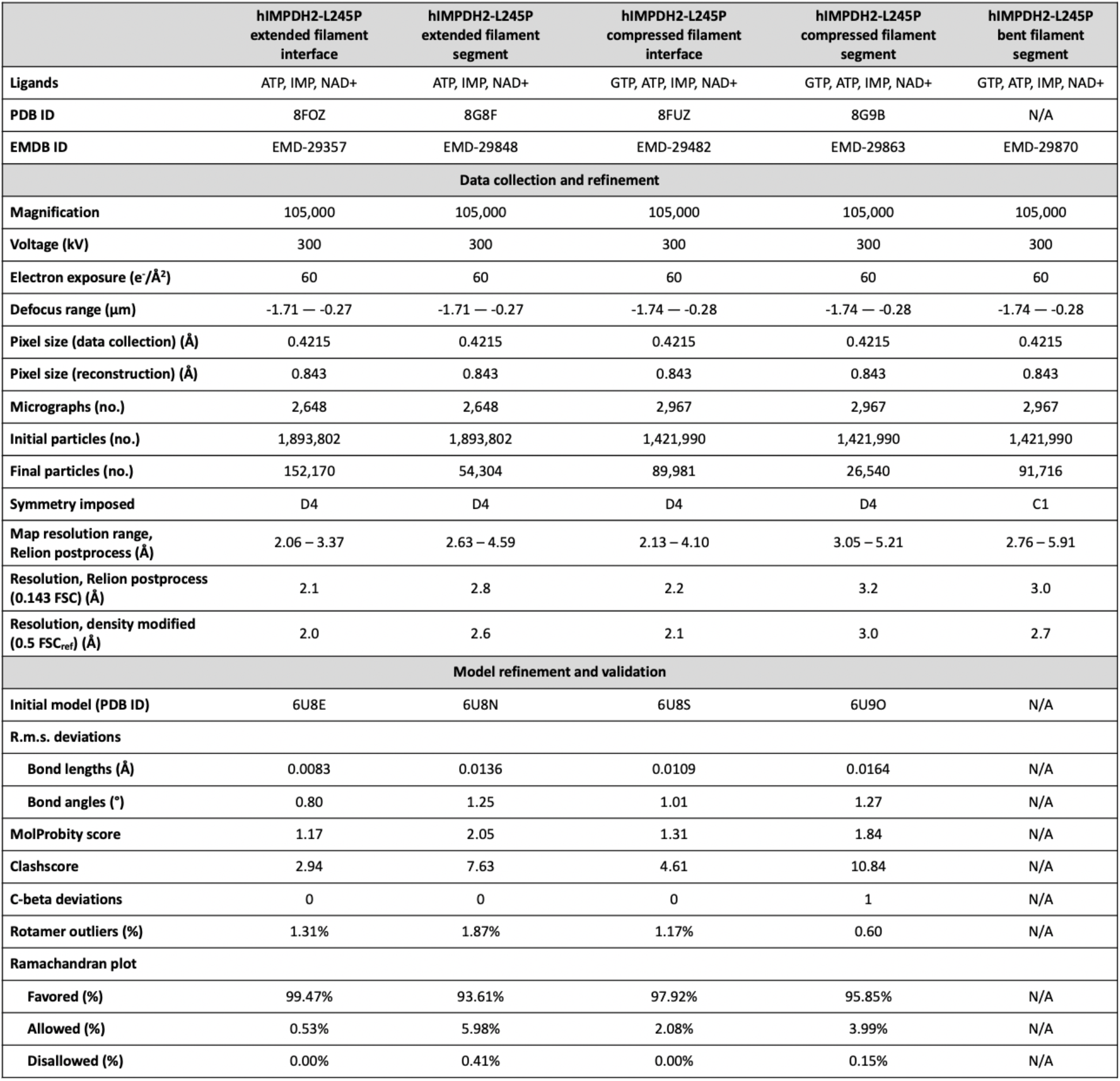
Cryo-EM data collection, refinement, and validation.

Particles from a subset of 100 micrographs were manually picked from each dataset, extracted, and classified in 2D to generate templates for autopicking. Particles were autopicked using these templates, extracted with a box size of 400 Å, with 4x binning, and classified in 2D to remove junk. For initial 3D refinement, a previous IMPDH filament reconstruction (EMD-20742 for the dataset with GTP, or EMD-20687 for the dataset without GTP) was low-pass filtered to 40 Å and used as an initial reference to align the particles at the filament assembly interface.

As described previously, two different locations along the D4-symmetric IMPDH filament may be used as symmetry origins–the center of the octameric filament segment or the center of the filament assembly interface (13, 14). Briefly, binned particles were initially aligned to a low-pass filtered IMPDH filament volume centered on the desired symmetry origin as a reference in an initial round of 3D auto-refinement, given a large offset range of 25. Helical segments were processed as single particles. Helical symmetry was not applied. After initial binned refinement, particles were un-binned and re-extracted with a pixel size of 0.843 Å, followed by 2D classification. Selected particles were submitted to 3D auto-refinement with D4 symmetry imposed. Remaining heterogeneity was removed with subsequent rounds of 3D classification.

CTF refinement and particle polishing were performed to refine per-particle defocus and per-micrograph astigmatism. Partial signal subtraction of the poorly resolved particle ends was performed on either the filament assembly interface or the octameric filament segment to boost resolution of the centered region. To this end, a mask was applied over either the central eight catalytic domains of the filament assembly interface or over the eight monomers of the octameric filament segment, and signal outside of the masks was subtracted. The resulting map was auto-refined in 3D. 2D and 3D classification with skip_align was performed to remove residual heterogeneity.

### Model building and refinement

Structures of human IMPDH2 filaments in the extended conformation (PDB 6U8N for the octamer-centered reconstruction and PDB 6U8E for the interface-centered reconstruction) and the compressed conformation (PDB 6U9O for the octamer-centered reconstruction and PDB 6U8S for the interface-centered reconstruction) were used as templates for model building. Templates were rigid-body fit into the cryo-EM maps using UCSF Chimera, and phenix.real_space_refine was used for automated fitting employing rigid-body refinement, NCS constraints, gradient-driven minimization and simulated annealing (70). Outputs from real-space refinement in PHENIX were inspected and manually adjusted with semi-automated fitting in ISOLDE and manual fitting in Coot (71, 72). This process was repeated iteratively, improving Molprobity statistics and fit to density. Refinement statistics are summarized in Supplemental Table 3. Structure figures were prepared using UCSF Chimera (73).

## Data availability

The coordinates are deposited in the Protein Data Bank with PDB IDs **8FOZ** (interface-centered extended hIMPDH2-L245P), **8G8F** (octamer-centered extended hIMPDH2-L245P), **8FUZ** (interface-centered compressed hIMPDH2-L245P), and **8G9B** (octamer-centered compressed hIMPDH2-L245P). The cryo-EM maps are deposited in the Electron Microscopy Data Bank with IDs **EMD-29357** (interface-centered extended hIMPDH2-L245P), **EMD-29848** (octamer-centered extended hIMPDH2-L245P), **EMD-29482** (interface-centered compressed hIMPDH2-L245P), **EMD-29863** (octamer-centered compressed hIMPDH2-L245P), and **EMD-29870** (octamer-centered bent hIMPDH2-L245P).

## Acknowledgements

We thank the patients and the family of the patients for participating in the study. We thank the Arnold and Mabel Beckman Cryo-EM Center at the University of Washington for electron microscope use. This work was supported by the US National Institutes of Health (R01GM118396 and S10OD023476 to J.M.K., T32GM008268 to A.G.O., and F31EY030732 to A.L.B.), and the German Research Foundation (DFG 458949627; ZE 1213/2-1 to M.Z.).

## Supplemental Notes

### Clinical report for case subject with L245P variant of IMPDH2

The proband was a 3-year-old female, the firstborn and only child of non-consanguineous parents of Jewish Ashkenazi origin. Pregnancy was significant for intrauterine growth retardation (IUGR), dilation of the lateral ventricles, and mild pulmonic stenosis on fetal ultrasound. Delivery was at 33+6 weeks, at a birthweight of 1520 grams (7th %ile) and with a head circumference of 28.5 cm (9th %ile). Complications of prematurity included bronchopulmonary dysplasia (BPD) with oxygen dependency for approximately one month. She was born with congenital dysplasia of the hip (CDH) which required surgical intervention, and with left sided torticollis. Echocardiogram revealed pulmonic stenosis, for which she underwent interventional catheterization (balloon angioplasty). There was also an atrial septal defect which closed spontaneously. Head ultrasound showed mild dilation of the left ventricle. Renal ultrasound was normal.

The proband had significant hypotonia and global developmental delay. She walked at around 2-years-7-months of age, and at 3 years of age, said less than 10 words, although she used nonverbal communication and seemed to have a higher receptive ability. Development quotient (DQ) was 42 at 14 months, and 60 at repeat evaluation. Hearing evaluation showed bilateral type B tympanograms and normal free field audiometry. She had abnormal posturing of the head, which she held tilted back and had a downward gaze. Ophthalmology exam including ultrasound of the orbits was normal. There were no convulsive episodes, and EEG was normal.

Family history was positive for CDH in the proband’s father and the paternal uncle. The mother’s half-brother was born with omphalocele and died at 2 years of age. His cognition was reported to be normal, and no DNA was available for testing. The maternal grandfather died in his 20’s of cancer affecting his leg; no further information was available.

On physical examination at 3-years-2-months, the child had failure to thrive, with a preserved head circumference of 47.5 cm (∼10th %ile). She was responsive, had good eye contact yet minimal facial mimicry. She held her head tilted back with abnormal posturing. She had a double hair whorl, sparse thin hair, high forehead, hypertelorism, bilateral epicanthal folds, a broad nasal bridge, open mouth, small low-set and posteriorly rotate ears, a systolic murmur, and axial hypotonia.

### Clinical report for case subject with K238R variant of IMPDH2

The proband was the product of a naturally conceived single gestation to a then 27-year-old G2P3 mother. Pregnancy and neonatal history were uncomplicated. Family history was non-contributory. He was born at 40+1 weeks gestation weighing 2920g (25th percentile) and was 51.4cm in length (75th-90th percentile). HC at birth was 33cm (25th percentile). APGAR scores were 8 at 1 minute and 9 at 5 minutes. He was discharged home at day of life 2.

Concerns for his development were first raised at 6.5 months of age due to gross motor delays. He sat at 8-9 months, crawled at 12 months, and walked at 18 months. Significant speech delay was also noted. Medical history was notable for tachypnea and stridor with feeds as a neonate, milk protein allergy and gastroesophageal reflux, meatal stenosis and penile adhesion s/p correction, easy bruising, torticollis and plagiocephaly, eczema, and hypotonia.

At 9 months of age, he weighed 8.396kg (24th percentile), was 74.8cm in length (80th percentile), and head circumference was 44.1cm (18 percentile). Physical examination was notable for slight upslant to palpebral fissures, prominent cheeks, supernumerary nipple, and mild hypotonia.

Follow-up at 23 months was notable for persistent developmental delay with most skills clustering around the 16-20 month age range per Developmental Pediatrician and concern for possible autism spectrum disorder. He was referred to Neurology for evaluation of possible movement disorders and was again noted to have hypotonia. He was started on Sinemet (6mg BID) due to the possibility of L-DOPA responsive symptoms in IMPDH2 related disorders with reported improvements in energy, interaction, speech skills, and movement.

## Supplemental Figures

**Supplemental Fig. 1: Photos of the patient with the K238R variant at different ages**.

[Images were removed for submission to bioRxiv]

Photos of the patient at 9 months (A), 16 months (B), and 26 months of age (C).

**Supplemental Fig. 2:**
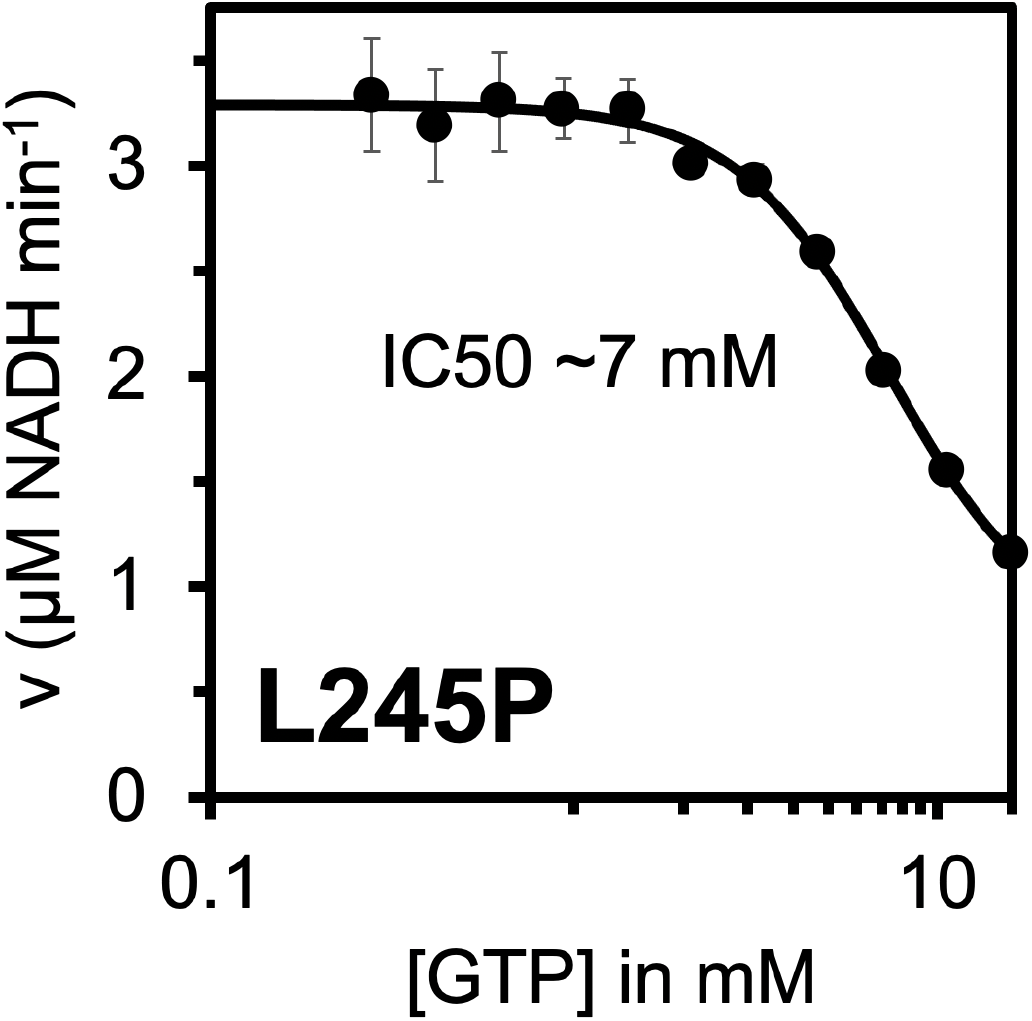
The L245P mutant is sensitive to high concentrations of GTP. GTP inhibition curve of the L245P variant up to 16 mM GTP. Each data point represents the average initial rate of three reactions. Error bars represent standard deviation for n=3 technical replicates. Velocities were calculated from the change in absorbance at 340 nm. Reactions were initiated with 300 μM NAD+ and contained 1 μM enzyme, 1 mM ATP, 1 mM IMP, 1 mM MgCl2 and varying concentrations of GTP.

**Supplemental Fig. 3:**
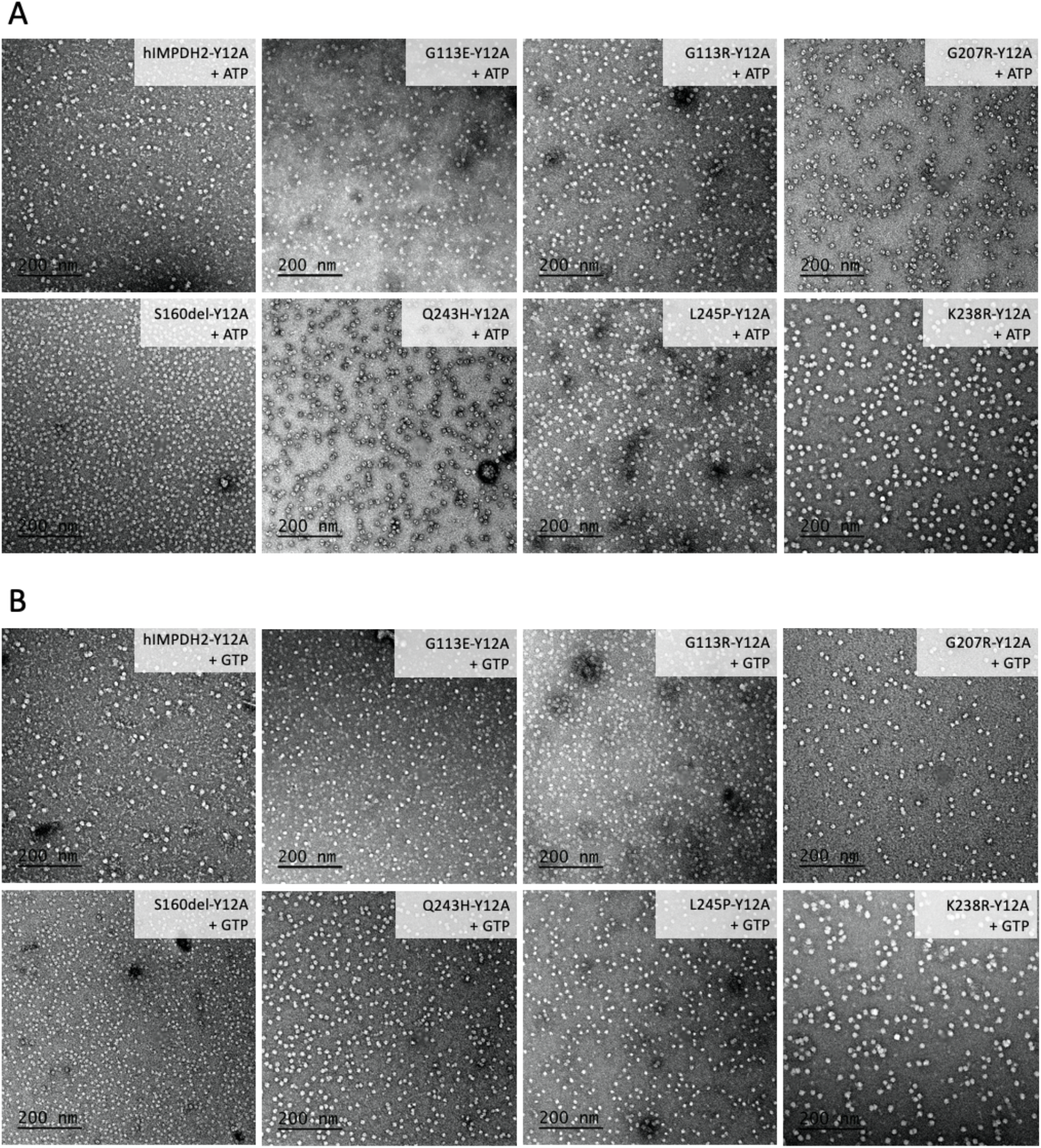
Non-assembly disease mutants imaged using negative stain EM. Representative negative stain images of 0.5 μM enzyme with either 1 mM ATP and 1 mM MgCl_2_ (A) or 5 mM GTP (B). Filaments were not observed in either condition.

**Supplemental Fig. 4:**
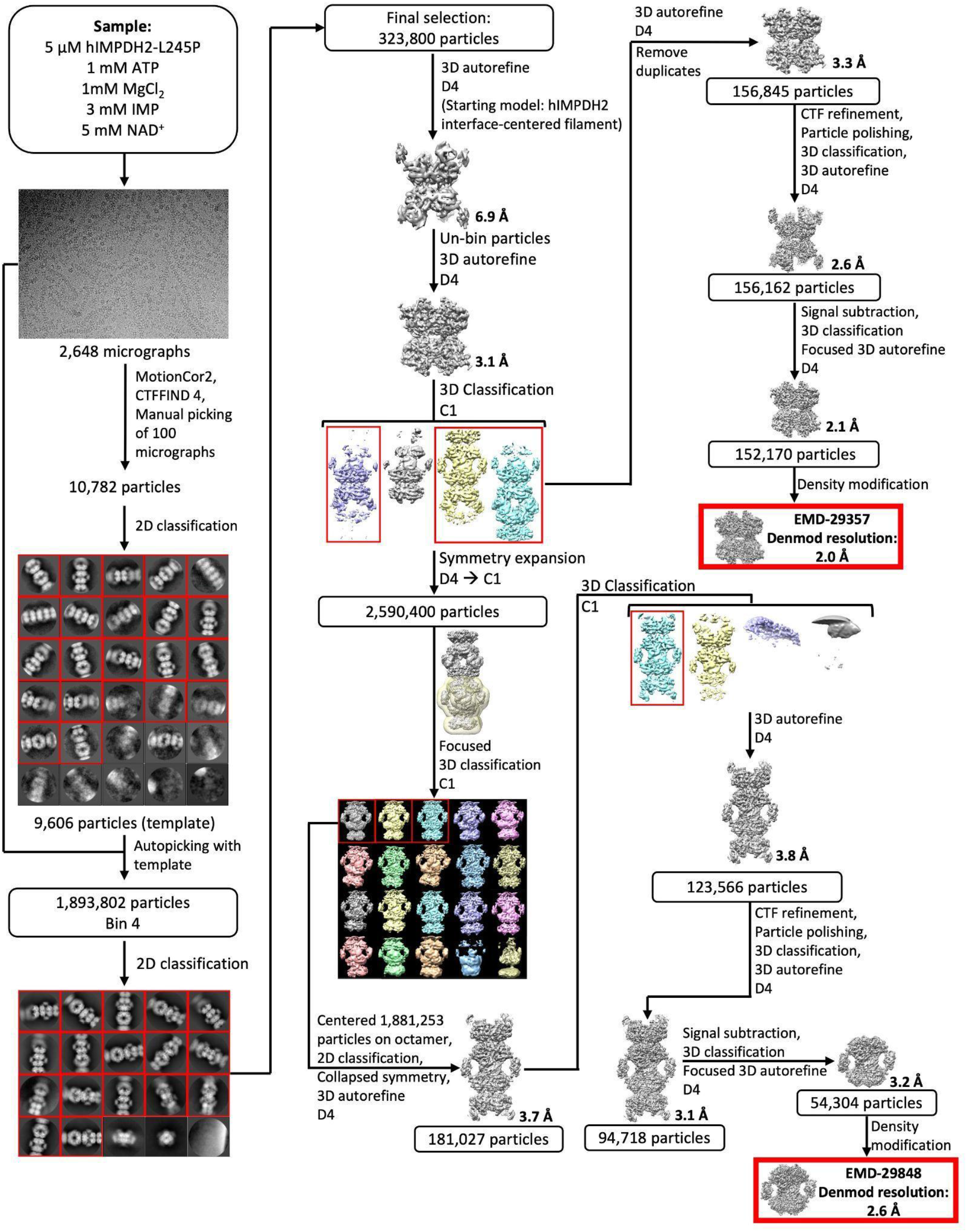
Extended L245P filament cryo-EM data processing.

**Supplemental Fig. 5:**
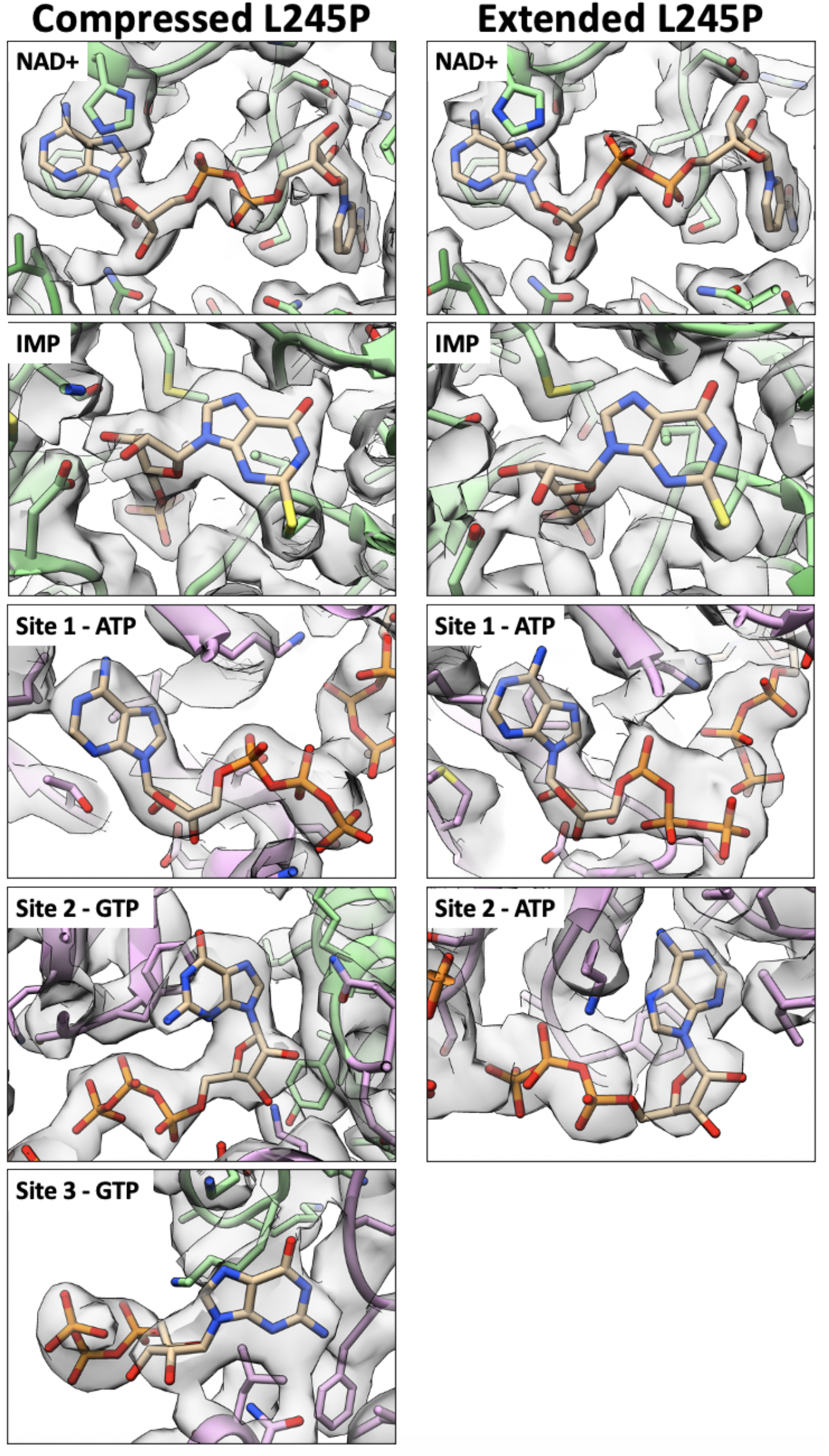
Volume around ligands in extended and compressed L245P structures. All ligands are resolved in the L245P extended and compressed structures. The catalytic domain is colored in green, and the regulatory domain is colored in pink.

**Supplemental Fig. 6:**
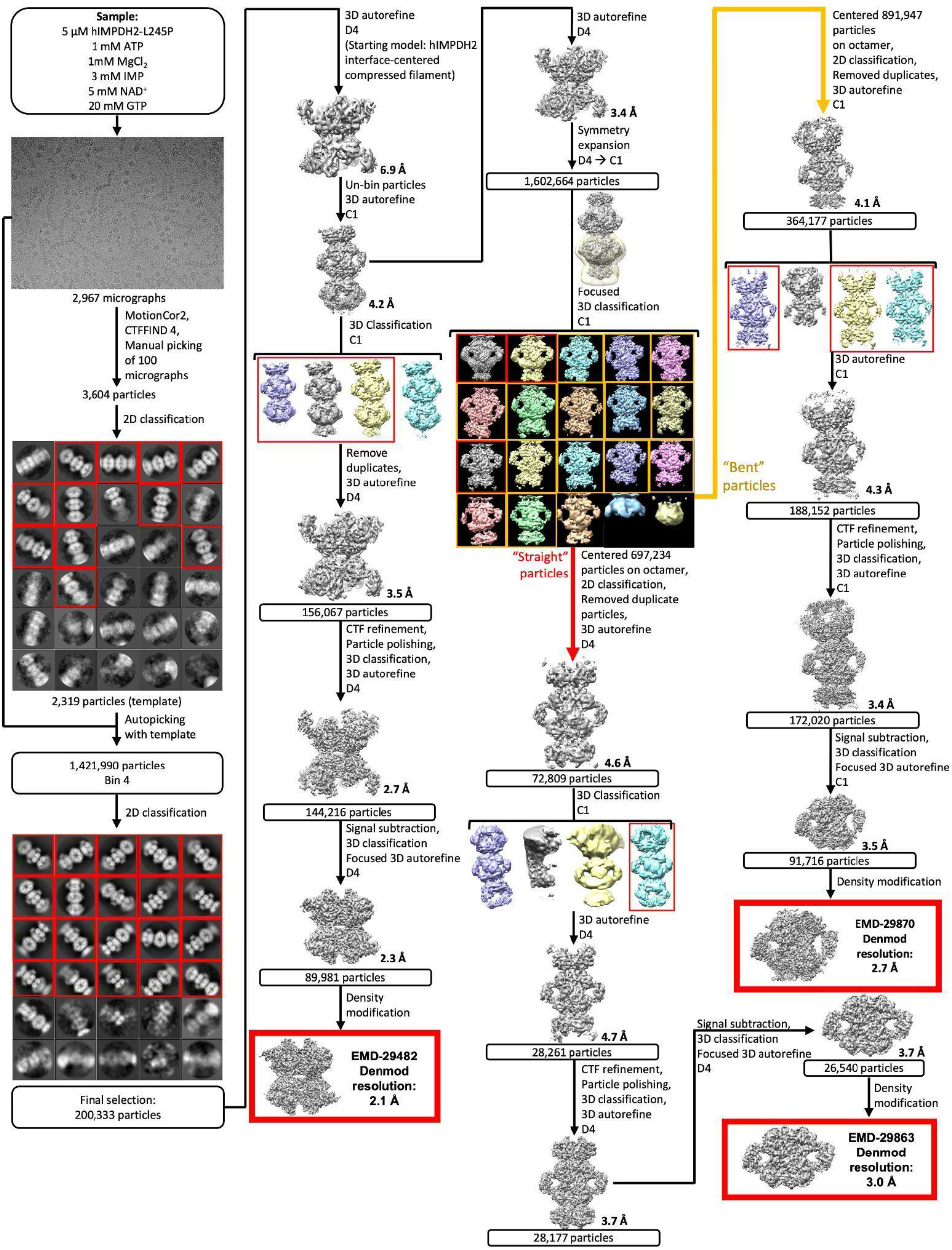
L245P+GTP/ATP/IMP/NAD+ data processing.

**Supplemental Fig. 7:**
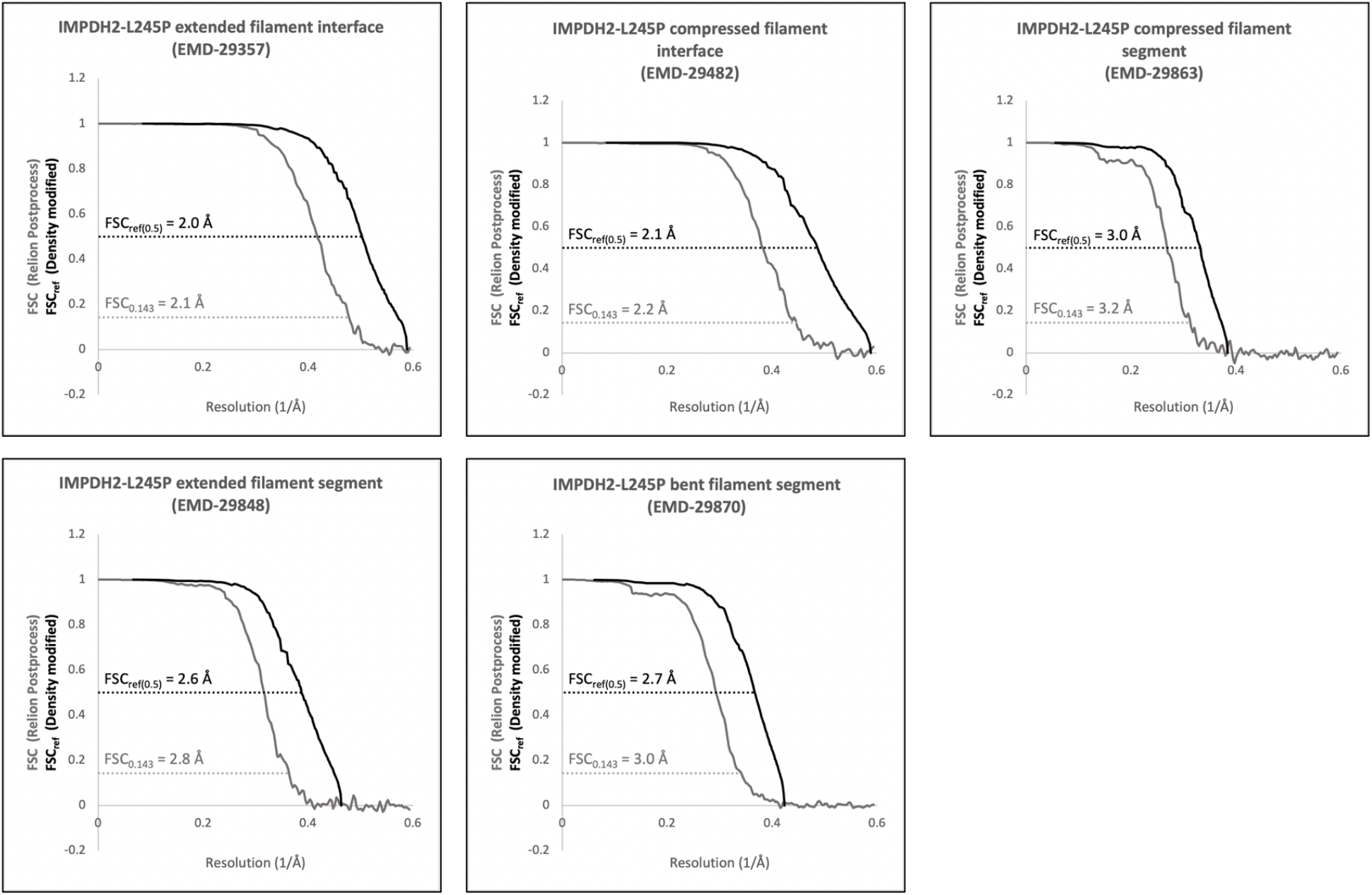
FSC curves of cryo-EM reconstructions.

**Supplemental Fig. 8:**
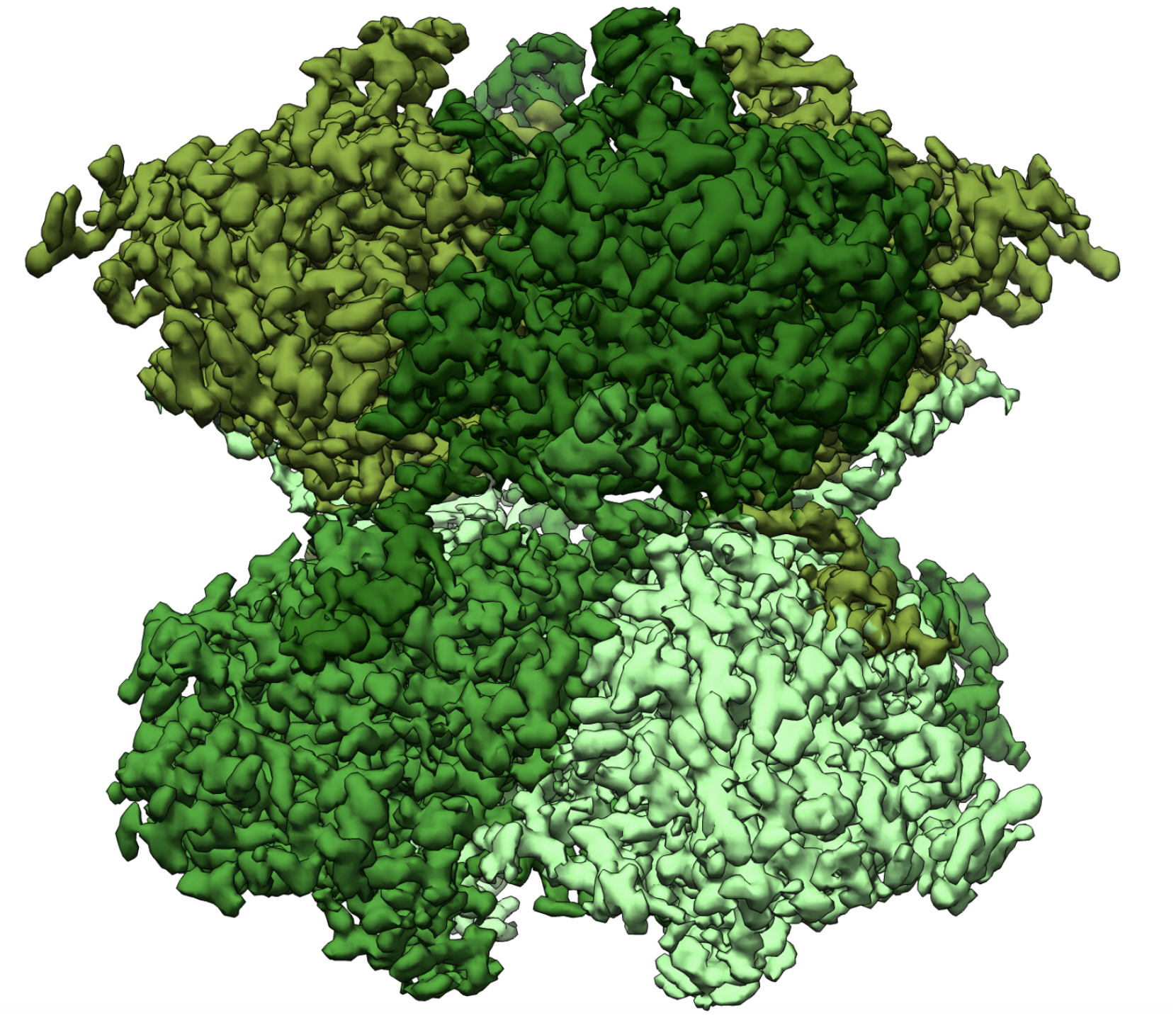
Filament assembly interface reconstruction of the L245P filament in the presence of 20 mM GTP, 1 mM ATP, 3 mM IMP, 5 mM NAD, and 1 mM MgCl2.

